# Motif selection enables efficient sequence-based classification of non-coding RNA

**DOI:** 10.1101/2023.09.19.558381

**Authors:** Ibrahim Chegrane, Nabil Benjaa, Aïda Ouangraoua

**Affiliations:** Département d’informatique, Université de Sherbrooke, Québec, Canada

## Abstract

**Motivation:** Non-coding RNAs (ncRNAs) classification is important for genome annotation and to perform functional analyses of these biological molecules. Efficient methods for large-scale RNA classification remain challenging. Existing methods often rely on structure similarity, and they require huge computing times because they use secondary structure information, which impedes any large-scale use.

**Results:** We present a sequence-based method that relies on the computation and the selection of common sequence motifs to provide a set of features for effectively classifying ncRNAs families by a supervised learning approach. The results show that our method achieves an equal or higher accuracy than existing structure-based methods and drastically reduces required computing times. Results also demonstrate that, thanks to an appropriate selection of local sequence motifs, an efficient sequence-based ncRNA classification can be achieved through supervised learning.

**Availability:** Code and datasets are available at https://github.com/chegrane/LSC-ncRNA

**Supplementary information:** Supplementary data are available at *Bioinformatics* online.

## 1 Introduction

Ribonucleic acids (RNAs) play important roles among living organisms. In Human, between 62% and 70% of the genome is transcribed into RNAs (Djebali et al., 2012), and about 98% of these RNAs are non-coding. Non-coding RNAs (ncRNAs) have various functions including gene expression control and guiding RNA modifications (Mattick, 2001). They are also involved in many human diseases such as neurological disorders and cardiovascular problems (Esteller, 2011). Depending on their function, ncRNAs are classified in various functional classes like ribosomal RNAs (rRNA), transfer RNAs (tRNA), microRNAs (miRNAs) and small interfering RNAs (siRNAs). Due to ncRNAs functional importance, there are many databases that provide information about ncRNAs. Most databases group ncRNAs into families characterized by sequence, structure, and functional similarities. Some general databases like RFAM (Kalvari et al., 2018) and RNAcentral (Consortium, 2017) contain various families of ncRNAs, while others are more specialized databases like miRBase (Kozomara et al., 2019) which only includes miRNA sequences. The widespread application of high-throughput RNA sequencing is generating increasingly large sets of new ncRNAs from various organisms. Some of these ncRNAs are members of established ncRNAs families and we need efficient classification methods to assign them to their families.

Several computational approaches exist for ncRNAs classification. Depending on the ncRNA families included in the classification, methods can be divided in two categories: 1) specific-purpose methods that aim to classify specific ncRNAs families (e.g., classification methods for tRNAs (Lowe and Eddy, 1997)), and 2) general-purpose methods to classify various ncRNAs family types. Depending on the information used for the classification, existing methods can also categorize as sequence-based or structure-based. Sequence-based classification relies solely on sequence similarity. Different techniques allow classifying ncRNAs based on sequence similarity. For instance, BLAST (Basic Local Alignment Search Tool) (Altschul et al., 1990) is an efficient heuristic alignment algorithm that is useful to find homologs of a given ncRNA sequence in a database of ncRNAs sequences. K-mer techniques rely on representing ncRNA sequences as vectors of k-mers counts to evaluate their similarity and classify ncRNAs (e.g., in (Kirk et al., 2018), a k-mer representation of ncRNAs was used for classification). The Profile Hidden Markov Models (profile HMMs) (Eddy, 1998) technique turns multiple sequence alignments of ncRNA family members into position-specific scoring systems, allowing to classify ncRNAs.

Structure-based classification uses RNA’s secondary structure information to improve ncRNAs classification accuracy. For instance, structure alignment techniques consist in aligning RNA secondary structures to evaluate their structural similarities and classify ncRNAs (e.g., in (Sanbonmatsu, 2016), RNA secondary structure alignment was used to classify long ncRNAs). Structural motif techniques consist in representing ncRNAs by vectors of structural features counts such as specific RNA structural elements or as measures of RNA folding attributes (e.g., in (Ng and Mishra, 2007), measures of global and intrinsic hairpin folding attributes were used to classify miRNAs). The Covariance Models (CM) technique builds stochastic context-free grammars (SCFGs) (Durbin et al., 1998) to represent RNA families based on multiple sequence alignments of RNA families with consensus structures. A CM is a generalization of a profile HMM that integrates secondary structure information. Recently, supervised learning techniques yielded promising results for both sequence-based and structure-based ncRNA classification. Some supervised learning-based methods such as (Noviello et al., 2020) use sequence information only, while other methods use secondary structure information, like (Ru et al., 2019). Most of these methods are specific-purpose methods, like circular RNAs (Chaabane et al., 2020) or lncRNAs and lincRNA (Amin et al., 2019), and most of them use a deep learning approach (Amin et al., 2019; Fiannaca et al., 2017).

Structure-based classification methods are generally more accurate than sequence-based ones. However, structure-based classification presents two main drawbacks. First, not all ncRNAs families have an established conserved secondary structure. Sometimes, the information about the structure is unavailable. Second, using the secondary structure information significantly increases ncRNAs classification computing times (Nawrocki and Eddy, 2013). To avoid these drawbacks, we need more accurate sequence-based classification methods for large-scale RNA classification. The superiority of structure-based classification, compared to sequence-based classification, usually stems from the higher conservation of ncRNA secondary structure compared to the primary sequence in several ncRNAs families, such as LncRNAs, or U2 and U4 spliceosomal RNAs. Nethertheless, even when members of an ncRNA family do not share globally conserved sequences, some sequence motifs are still locally conserved and can help identify family members. In this paper, we follow this path and present a general-purpose sequence-based ncRNA classification method that relies on the computation and the selection of conserved sequence motifs, and a supervised learning approach.

Our method is divided into two main steps. The first step is the computation and the selection of conserved sequence motifs for the set of ncRNA families included in the classification. The second step consists in selecting a set of classification algorithms that allows achieving the highest accuracy in a reasonable time. We experimentally compared our method with existing sequence- and structure-based methods relying on various techniques, which shows that the proposed method achieves a comparable or a higher accuracy than existing methods, while drastically reducing the computing time. In the case of low sequence conservation within ncRNA classes, our sequence-based method outperforms structure-based methods.

The remaining of the paper is organized as follows. Section 2 describes the two steps of the method and the training dataset. Section 3 presents the results of the experimental analyses to set parameters values for motif selection and to select the best classification algorithms. We also present and discuss the results of the experimental comparison of our method with existing methods. Finally, section 4 concludes.

## 2 Methods

The method is divided into two steps. The first step is the computation and the selection of sequence motifs that allow defining a vectorial representation of ncRNAs sequences. The second step is the selection of supervised learning classification algorithms that allow achieving the most accurate classification of ncRNA sequences. For both steps, the selection process was realized by searching the motif selection parameter space and the classification algorithm space for the best cross-validation settings.

The performance measures used to compare different settings regard the size of the data generated, the data generation time, the training time, the classification time, and the classification accuracy. The classification accuracy is the rate of correct classifications. Let *y*_i_ and *ŷ*_i_ be the real and predicted class for the i-th RNA sequence (*i ∈* [1 − *n*]), then the accuracy is defined as 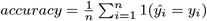.

### 2.1 Step 1: Sequence motifs computation and selection

Aside from sharing a conserved secondary structure, members of an ncRNA family often display local conserved sequence motifs even when the overall sequence similarity is low. These motifs can be used as features to represent ncRNA sequences for classification. We hypothesize that the discriminating power of these common sequence motifs is greater than that of k-mer features, traditionally used for representing and classifying ncRNAs through supervised learning approaches. In this step, the method computes all sequence motifs that appear in at least two members of an ncRNA family. Next, the initial set of sequence motifs is filtered according to various tuned parameters, and a final subset of selected motifs is used to represent ncRNA sequences for classification.

#### 2.1.1 Common motifs computation

Given an ncRNA family, a common motif is a substring that appears in at least two family members. To compute the set of common motifs for all ncRNA families, the method uses a generalized suffix tree (GST) (Bieganski et al., 1994; Gusfield, 1997). A GST is a suffix tree constructed over multiple sequences. Given a set of RNA sequences, a GST can be constructed in a linear time using the Ukkonen’s algorithm (Ukkonen, 1995). An example of GST for 4 sequences appear in Supplementary Figure 9. Given a set of *m* ncRNA families *{f*_1_, *f*_2_, …, *f*_m_*}* such that each family *f*_j_ contains |*f*_j_ | sequences, and the total number of sequences is 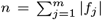, a GST is built for all sequences of all *m* families. Building one GST for all sequences of all families, instead of building one GST for each ncRNA family, allows reducing the size of the memory space required to store the GST, because sequences of different families share several substrings. It also allows computing a set of unique common motifs directly for all families, thus avoiding an additional processing time to compute the union of all family common motif sets.

This GST allows finding the locations of all the occurrences of a substring in all sequences. Simply traverse the GST, and collect, at each node, the corresponding substring with the list of leaf labels of the tree rooted at this node. From the list of leaf labels associated to a substring, the number of substring occurrences in each sequence is computed and only substrings with occurrences in at least two members of a family are kept as common motifs. The maximum length of a common motif is limited to 20 nucleotides, because most common motifs are small, and limiting the maximum size of common motifs reduces the GST processing time for large datasets. The common motifs with their numbers of occurrences are stored in a *n* × *p* matrix *M* whose lines correspond to the *n* RNA sequences, and whose columns correspond to the set of *p* sequence motifs, such that any cell *M* [*s*_i_][*cm*_k_] contains the common motif number of occurrences *cm*_k_ in sequence *s*_i_.

#### 2.1.2 Common motifs selection

Motif selection allows reducing the dimension of the RNA sequence representation by selecting the most discriminating common motifs for ncRNA classification. This crucial step allows achieving an efficient and effective classification. Thus, to reduce the computing time of the classification while preserving the classification high accuracy, we applied various filters to remove redundant information and to select the most informative motifs. The filters parameters are described below, and in section 3, we describe and discuss the results of the cross-validation process to find the best values of filtering parameters.

##### A. Filtering out submotifs (parameter *submotif*)

The GST processing allows retrieving all common motifs of ncRNA families. Some of the collected motifs are substrings of other common motifs and provide redundant information. A motif *x* is an exact submotif of a motif *y* if, and only if, *x* is a substring of *y*, and *x* and *y* are substrings of the same ncRNA sequences with the same number of occurrences in each sequence.

To filter out exact submotifs from the set of common motifs, the set of acceptable motifs is initialized to an empty list *L* of motifs sorted by increasing length. Next, for each new common motif *x*, if *x* is not an exact submotif of a longer motif already appearing in *L*, and there is no shorter motif in *L* that is an exact submotif of *x, x* is added in *L*. The method considers two values for the parameter *submotif* : F if submotif are filtered, and NF otherwise.

##### B. Filtering according to motif length (parameters *minl* and *maxl*)

In theory, the size of motifs common to two sequences may range from 1 nucleotide to the length of the shortest sequence. The maximum size is reached when we have identical sequences or when one sequence is a substring of the other. However, in practice for ncRNA families, it rarely happens, and common motifs usually have limited lengths. Hence, the method considers two parameters that limit the size of selected common motifs: *minl* and *maxl* are the minimum and maximum length parameters used to filter common motifs.

##### C. Filtering according to motif conservation in their family (parameter *α*)

Given a common motif of an ncRNA family, the representative power of the motif in the family increases when the number of sequences containing the motif increases. A common motif that occurs in all sequences of the family is more representative of the family than a common motif present in only two sequences. Let *x* be a common motif that occurs in *k* out of *n* sequences of an RNA family. The representative power of *x* in the family is 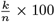. The method considers a parameter *α* whose values range between 0 and 100 as the minimum representative power required for a common motif to be selected. For instance, if a family includes 100 sequences and *α* = 20, a common motif will be selected if, and only if, it occurs in at least 20 sequences.

##### D. Filtering according to the variation of the number of occurrences (parameter *β*)

The variation of the number of occurrences of a common motif in a family sequences also indicates the motif conservation in the family. A common motif that has the same number of occurrences in all sequences where it appears is better conserved than a common motif whose number of occurrences varies between sequences. Let *x* be a common motif whose lowest number of occurrences in a sequence where it appears is *l* while its highest number of occurrences in a sequence is *h*. The occurrence difference of *x* in the family is the difference *h*−*l*. The method considers a parameter *β* as the maximum occurrence difference required for selecting a common motif. For instance, if a family contains 3 sequences and *β* = 2, a common motif that has a number of occurrences 2, 3, 5 will be discarded, while a common motif that has a number of occurrences 2, 3, 4 will be selected because its occurrence difference is 4 − 2 = 2, which meets the upper bound *β* = 2.

##### E. Filtering according to the number of common motifs occurrences (parameter *γ*)

Besides parameters *α* and *β* that allow considering various levels of common motifs conservation in their families, we also considered the number of common motif occurrences in a sequence. The method considers an additional parameter *γ* as the motif minimum number of occurrences to be selected. Let *x* be a common motif whose minimum number of occurrences in any sequence where it appears is *occ*(*x*). The motif *x* is selected if, and only if, *occ*(*x*) *≥ γ*. For instance, if a family contains 3 sequences and *γ* = 2, a common motif *x* that has a number of occurrences 1, 2, 3 in the 3 sequences will be discarded because *occ*(*x*) = 1 which is lower than *γ*.

### 2.2 Step 2: Classification algorithms selection

Several supervised learning approaches exist for classification (Kotsiantis et al., 2007; Singh et al., 2016) from which we aim to choose the best algorithms based on their ability to efficiently classify large numbers of ncRNA families. The data for the classification are represented using cardinal features. Based on the type of input data, we preselected seven (7) classification algorithms able to handle this type of data (Kotsiantis et al., 2007): Decision Tree (DT) (Murthy, 1998); Random forest (rdf) (Breiman, 2001); ExtraTree (EXT) (Geurts et al., 2006); Support vector machine (SVM) (Cortes and Vapnik, 1995); K-nearest neighbors (KNN) (Cover and Hart, 1967); Gaussian Naive Bayes (GNB) (John and Langley, 2013); and MultiLayer Perceptron (Neural networks) (MLP) (Rosenblatt, 1961; Taud and Mas, 2018). Next, based on experimental comparison, we selected the best algorithms in terms of classification time and accuracy. The comparison also includes a voting model based on the selected algorithms.

### 2.3 Training dataset and experimental design

We used data from the RFAM database (called Dataset #1) to select common sequence motifs and classification algorithms, and to train and test our classification methods. RFAM describes ncRNA families (Kalvari et al., 2018). For each family, the seed alignment is an alignment of a manually curated sample of family representative sequences called seed sequences. The RFAM database also defines ncRNA clans, which are a meta classification to group ncRNA families for which homology can be recognized, but the sequence similarity is too low to yield good alignments (Gardner et al., 2010). We used two additional datasets from (Noviello et al., 2020) with noisy sequences (called Dataset #2) and from RFAM on ncRNA clans (called Dataset #3) to compare the performance of our method with existing classification methods.

#### 2.3.1 Dataset #1

The RFAM database version 14.1 includes all ncRNAs families that contain at least 4 different seed sequences: 2,246 ncRNAs families with a total of 70,122 seed sequences. The dataset is imbalanced and includes 1, 064 families of size in the interval [4, 10], 500 families for [11, 20], 364 for [21, 40], 191 for [41, 100], 119 for [101, 500], 4 for [501, 1000], and finally 4 for [1001, 1542]. Each dataset class was split into two random subsets for training (70%) and testing (30%).

The training dataset was used to search the motif selection parameter space and the classification algorithm space by cross-validation. However, exhaustively considering all values combinations of parameters and models requires huge computing time and memory space. Hence, we designed a faster search strategy that relies on two aspects.

First, we evaluated candidate values of parameters on a small portion of the training dataset. We considered two (2) samples of the training dataset that preserved the proportion of ncRNA families in each of the size groups [4, 10], [11, 20], [21, 40], …, [1001, 1542]: a small dataset including 350 ncRNA families for which we considered subsamples of increasing sizes {50,150,250,350}; a medium dataset including 600 ncRNA families for which we considered subsamples of increasing sizes {100,200,300,400,500,600}.

Note that each subsample of the small and medium datasets also preserved the proportion of ncRNA families in each of the size groups to reflect the imbalanced property of the dataset.

Second, the search strategy followed an iterative selection process such that parameters were included iteratively in this order *submotif, minl*_*maxl, α, β, γ, classification*_*algorithm*. At each iteration, only the best candidate values of parameters were selected for the next iteration. Each iteration consisted in a stratified 10-fold cross-validation that compared all combinations of values of parameters considered in the step to select the best candidates for the next step. For the tuning of parameter sets including *minl*_*maxl, α, β, γ*, we only used two classification algorithms: Extra-tree (EXT) and neural networks (MLP), because they achieved the best classifications regarding the accuracy among all classification algorithms on the 100-family sample dataset. Note that the superiority of EXT and MLP is confirmed in section 3.2 in the experiments for the selection of classification algorithms.

After completing the selection of common motifs for RNA representation and of classification algorithms, we trained the models on the whole training dataset and tested them on the test dataset. Besides the classification accuracy, we also used the F-beta score to compare methods. It is defined as follows. The precision evaluates the ability of the model not to predict a class for an RNA sequence if it does not belong in this class. For each class, *Precision* = *T P/*(*T P* + *FP*), where TP and FP are the numbers of True Positives and False Positives. The recall evaluates the ability of the model to predict the real class for all class members. For each class, *Recall* = *T P/*(*T P* + *FN*), where TP and FN are the numbers of True Positives and False Negatives. For each class, the F-beta score is the harmonic mean of the precision and recall. For each classification, the F-beta score is a mean of all classes F-beta scores.

#### 2.3.2 Dataset #2

The benchmark dataset provided in deepncRNA (Noviello et al., 2020) allowed comparing our method with existing methods. It includes 306,016 sequences distributed among 88 RFAM families of small ncRNA such that each family contains more than 400 sequences. This dataset is available at https://github.com/bioinformatics-sannio/ncrna-deep. Each dataset class was split into three random subsets for train (84%), validation (8%), and test (8%) in (Noviello et al., 2020). We kept the initial partition of the dataset to be able to compare our results with those of (Noviello et al., 2020). Since no validation is performed on this dataset, we merged the train and validation sets into a training dataset representing 92% of the data.

Based on this dataset, we generated a noise dataset as follows. For each sequence in the test dataset, we added x% noise at its start and its end, i.e., x/2% at each extremity, such that the percentage x% of noise compared to the length of the original sequence varied between 0% and 200% with steps of 25%. The noise was generated by randomly shuffling the sequence while preserving the nucleotide and dinucleotide frequency of the original sequence. This dataset allowed evaluating the robustness of the classification methods noise-wise, as real-life applications to classify new unknown RNA sequences. We trained the compared methods on the training dataset and tested them on the test dataset with various percentages of added noise. We compared the methods based on the training time, the classification time and the classification accuracy.

#### 2.3.3 Dataset #3

From the RFAM database, we build a dataset to evaluate the performance of the classification methods for ncRNA classes with low sequence similarity versus high sequence similarity. The low sequence similarity dataset included 36 classes defined as the clans of RFAM 14.8 containing at least 4 ncRNA families. This resulted in 36 clans containing 4 to 11 families per clan and a total of 199 families for all 36 clans. For each clan, we selected 12 sequences from each family. The high sequence similarity dataset included 199 classes defined as the 199 ncRNA families contained in the 36 clans from the low sequence similarity dataset. Each class of each dataset was split into two random subsets for train (70%) and test (30%). We trained and tested the compared methods on both the low and the high sequence similarity datasets. We compared the methods based on the classification accuracy.

### 2.4 Implementation

The code for the method is written in C++ (STANDARD 14) for the computation of common motifs, and in Python 3.8 for the selection of common motifs and classification algorithms and the comparison with existing methods. The source code is available under LGPL license at https://github.com/chegrane/LSC-ncRNA. All experiments were performed on a computer server model Intel(R) Xeon(R) CPU E5-2660 v4 @2.00GHz, with 48 CPUs and 1024 Go of memory, running under Linux operating system (Ubuntu 16.04.6 LTS).

## 3 Results

In this section, we discuss the results of the experimental analyses conducted to choose the best values of parameters for motif selection and the best classification algorithms to achieve an efficient classification of large ncRNAs sequences datasets. We also discuss the results of the compared methods.

### 3.1 Computation and selection of sequence motifs

#### Computation of the initial set of motifs

The initial set of motifs consists of all common linear motifs (substrings) of length included between *minl* = 2 and *maxl* = 20 between any two sequences of a family. Figure 1 (Left) shows, in solid line, the size evolution of the resulting datasets for a number of families 100 to 600 in the medium dataset. We see that the dataset size increases quickly along with the number of families increasing. For 100 ncRNAs families, the generated dataset is 1.3 Go, while it increases to 120 Go for 600 families. Therefore, to efficiently classify large datasets like the RFAM dataset, we must perform a selection of motifs to reduce the dimension of the dataset.

**Fig. 1.**
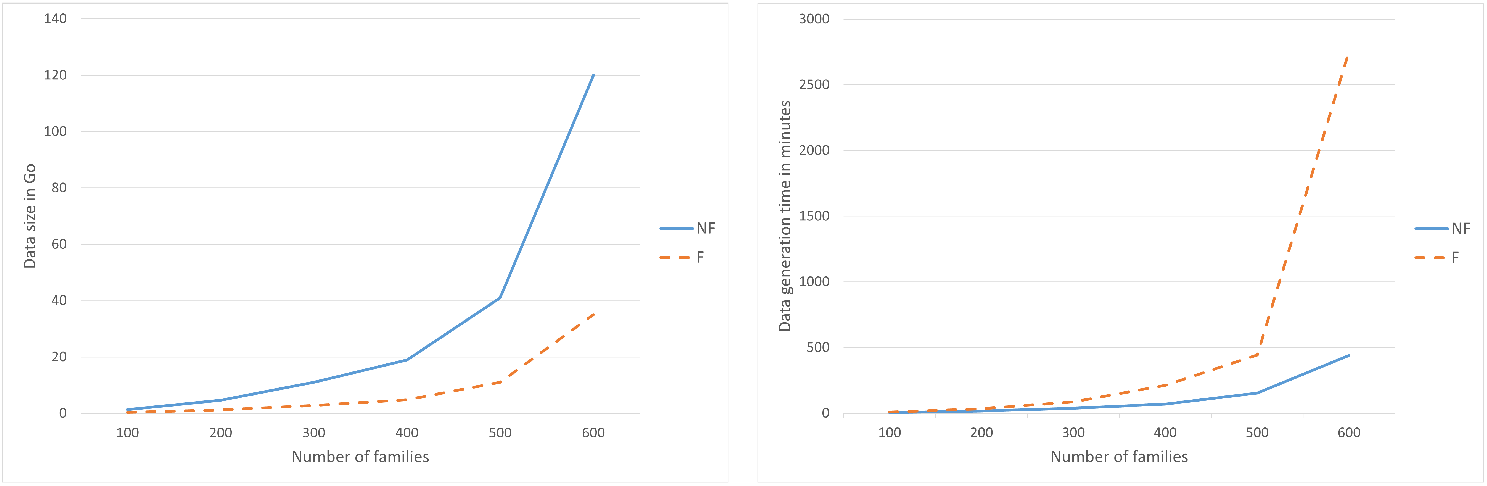
Relation between the number of families and: (Left figure) the data size; (Right figure) the data generation time. The relations are displayed for the generation of dataset with *submotif* = F and *submotif* = NF.

#### A. Filtering out submotifs (parameters *submotif*)

In this experiment, all exact submotifs are filtered from the initial set of motifs. Figure 1 (Left) shows, in dashed line, the size evolution of the resulting datasets for a number of families 100 to 600 in the medium dataset. Figure 1 (Right) shows the evolution of the computing time in minutes for the initial set of motifs, and for the computation including the filtering of exact submotifs. Removing the exact submotifs considerably reduces the size of the dataset. Nevertheless, the filtering process adds a considerable computing time. It requires about 2, 3 and 6 times more computing time for a number of families 100, [200 − 500], and 600 respectively. We also see in the sequel, in combination with the parameter *α*, that deleting the exact submotifs does not impact the classification accuracy (see Figure 3 (Right)). Therefore, in the next steps, submotifs were not filtered (i.e., *submotif* = NF).

#### B. Filtering according to motif length (parameters *minl* and *maxl*)

In this experiment, we compared settings where the size of the initial set of motifs was reduced by limiting the size of the common motifs using parameters *minl* and *maxl*. We considered two collections of settings: 20 fixed-length settings defined by *minl* = *maxl* = *k* for *k ∈* [1..20], and 18 combined-length settings defined by *minl* = 2 *maxl* = *k* for *k ∈* [3..20]. Figure 2 and Supplementary Figure 10 show the evolution of the number of motifs (Top) and the classification accuracy (Bottom) for the small dataset ncRNA families. Regarding the fixed-length settings, we see that the EXT classification algorithm reaches the highest accuracy (*>* 0.95) for all numbers of families (50, 150, 250, 350) at *minl* = *maxl* = 5, and the MLP algorithm at *minl* = *maxl* = 7. For both classifiers, the accuracy decreases with the number of families increase. Moreover, up to *minl* = *maxl* = 7, the number of motifs is less than 20*K* for all numbers of families, then, it increases quickly to reach a pic at *minl* = *maxl* = 10. After *minl* = *maxl* = 11, the number of motifs decreases because long motifs are less conserved in sequences. Regarding the combined length settings, we see that the EXT classifier reaches the highest accuracy 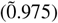 at *maxl* = 5, and the MLP classifier at *maxl* = 9. Compared to the accuracy achieved using fixed length motifs, the combined length motifs allow achieving a higher accuracy. The number of motifs for combined length increases quickly, especially after *maxl* = 8. Note that for both fixed length and combined length, we see that the EXT algorithm performs better with small length motifs compared to MLP. Indeed, EXT handles small size datasets better, as it relies on a decision tree approach, whereas the MLP methods rely on a neural networks approach which requires large-size datasets. Also note that for both fixed length and combined length, the processing time is correlated to the number of motifs (data not shown). Based on these results, in the next steps, fixed length parameters were limited to *minl* = *maxl* = *{*5, 6, 7*}*, and combined length parameters to *minl* = 2 and = *maxl* = *{*5, 6, 7, 8*}*. The maximum inclusion-wise dataset defined by *minl* = 2 and *maxl* = 8 was used in steps C to E. Note that the lower bound *minl* is less important than the upper bound *maxl*, as the number of small length motifs is very small compared to long length motifs.

**Fig. 2.**
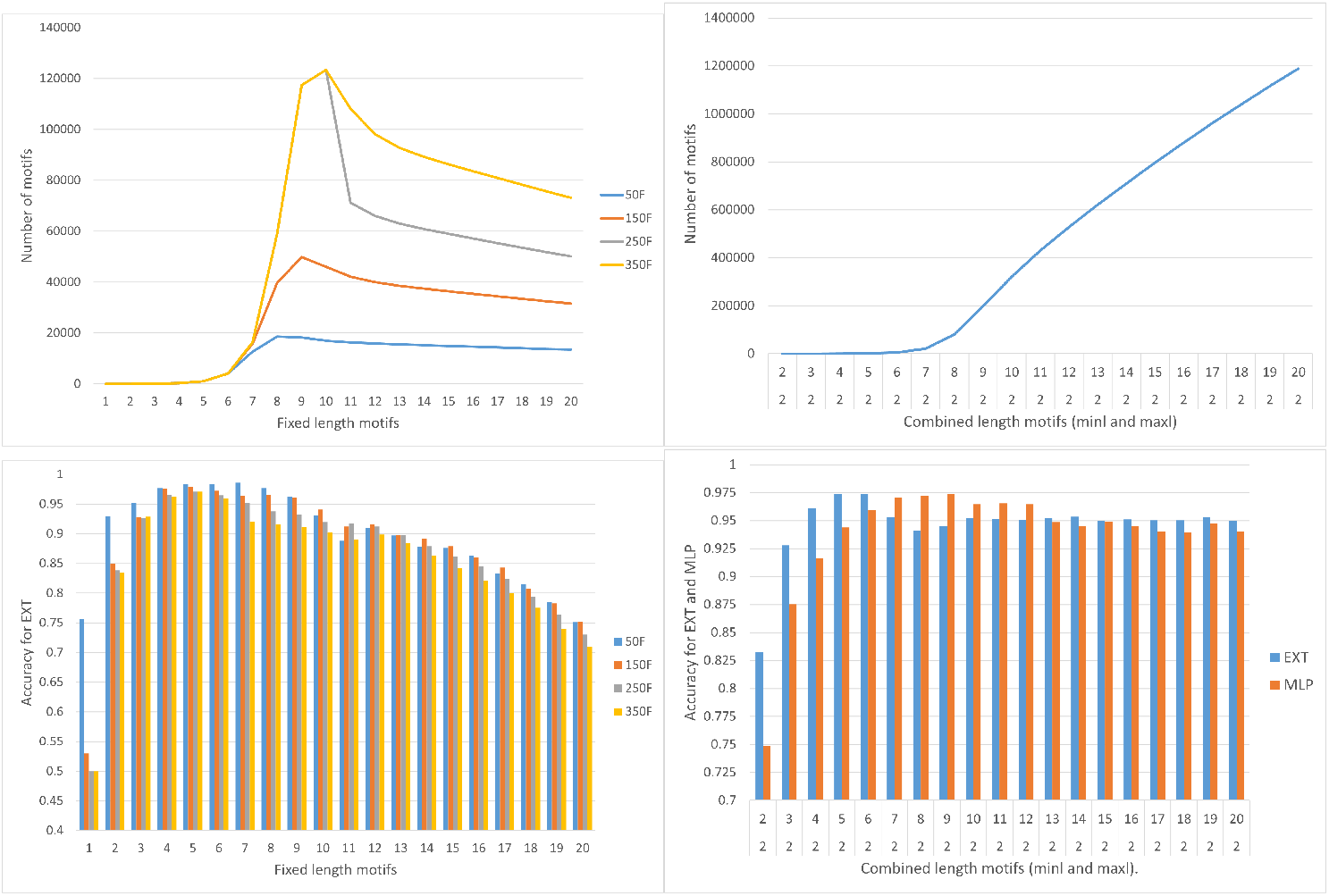
Relation between the number of generated motifs and : (Top-Left figure) the size of fixed-length motifs; (Top-Right figure) the size of combined-length motifs. Classification accuracy for: (Bottom-Left figure) different sizes of fixed-length motifs using EXT; (Bottom-Right figure) different sizes of combined-length motifs using EXT and MLP.

**Fig. 3.**
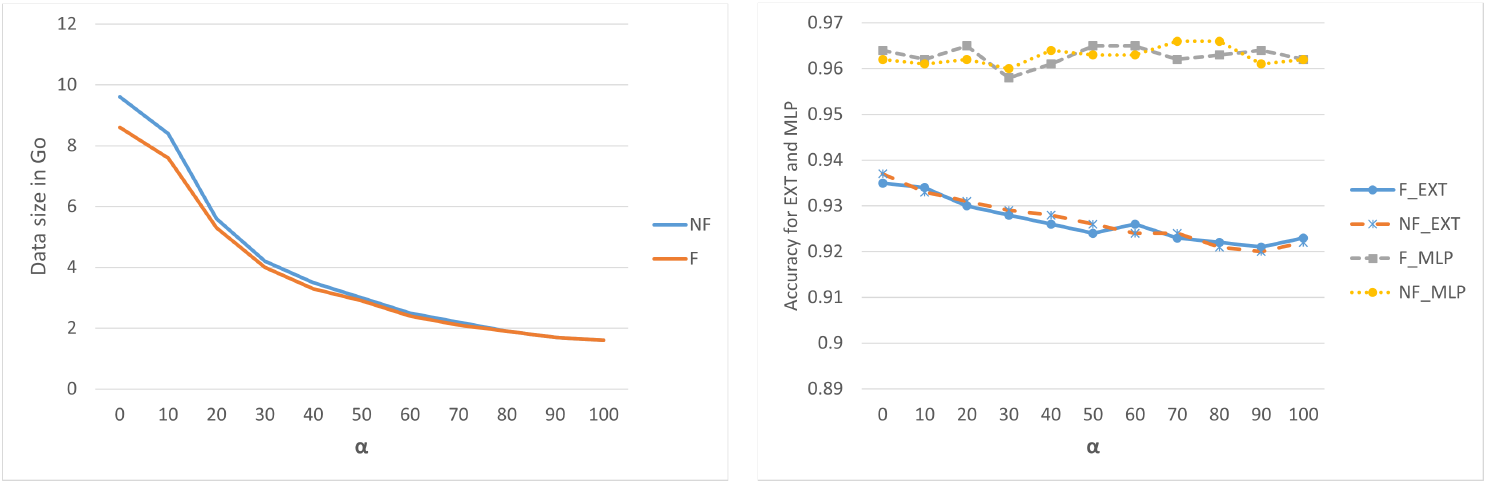
Relation between *α* and : (Left figure) the data size; (Right figure) the classification accuracy for EXT and MLP with *submotif* = F or *submotif* = NF.

#### C. Filtering according to motif conservation in their family (parameter *α*)

In this experiment, we compared different settings including the parameter *α*. Figure 3 and Supplementary Figure 11 show the evolution of the dataset size, the classification accuracy, the training time and the data generation time for different values of *α*. The figures illustrates that using *α* effectively reduces the size of the dataset. It also allows to reduce the proportion of exact submotifs in the dataset, because long motifs are less conserved between sequences. Therefore, using *α* results in discarding several long motifs which are likely to contain shorter exact submotifs. Moreover, using the parameter *α* or the parameter *submotif* has very little effect on the classification accuracy. These results indicate that filtering exact submotifs is useless when using *α*, and using *α* allows reducing the dataset size and the computing time required while preserving the classification accuracy. Therefore, in the next steps, the comparisons included values *{*0, 50, 100*}* of *α*.

#### D. Filtering according to the number of occurrences variation (parameter *β*)

We then compared different settings including the parameter *beta*. Supplementary Figure 12 compares the number of motifs for *β* = 0 and *β* = +*∞* for different values of *α*. Surprisingly, for all values of *α*, the number of motifs remains almost the same with *β* = 0 and *β* = +*∞*. After investigation, we found out that for most motifs, the number of occurrences in sequences is the same and equals 1. This explains why *β* = 0 impacts the filtering very little. Therefore, we discarded the *β* parameter in the next steps.

#### E. Filtering according to the number of occurrences per sequence (parameter *γ*)

In this experiment, we compared different settings including the parameter *gamma*. Figure 4 and Supplementary Figure 13 compare the classification accuracy and the number of motifs for *γ* = 1 and *γ* = 0 for different values of *α*. Surprisingly, the accuracy of the classification remains comparable for *γ* = 1 and *γ* = 2 with the two classifiers EXT and MLP. EXT yields a slightly higher accuracy for *γ* = 2, while MLP shows the opposite. As expected, because most common motifs have a single occurrence in sequences, the number of common motifs for *γ* = 2 is drastically reduced compared to *γ* = 1. In the next step, the comparisons included values *{*1, 2*}* of *γ*.

**Fig. 4.**
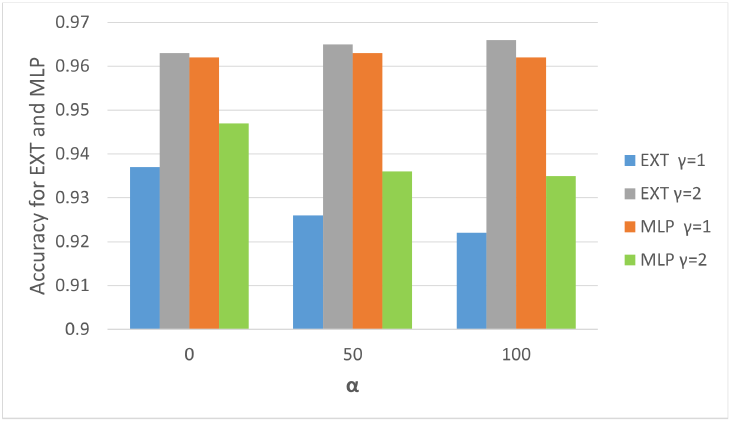
Relation between *α* and the classification accuracy for EXT and MLP with *γ* = 1 or *γ* = 2.

### 3.2 Selection of a classification methods

This section compared the classification algorithms performances. We considered the combinations of the following selected settings: either fixed length motifs with *minl* = *maxl ∈ {*5, 7*}* (denoted *F* = 5 and *F* = 7), or combined length motifs with *minl* = 2 and *maxl ∈ {*5, 8*}* (denoted *C*[2, 5] and *C*[2, 8]); *α ∈ {*0, 50, 100*}*; *γ ∈ {*1, 2*}*. Figure 5 shows the results for *α* = 50, which yields the best compromise between classification accuracy and computing time for all settings. We see that the three algorithms EXT, RDF, and MLP outperform the other methods in terms of accuracy with reasonable computing times in all parameter settings. Therefore, we also included in the comparison a combination of the three methods into a voting model with voting scores (1,1,1). The voting model increased the accuracy of the classification compared to the individual models, but it naturally requires longer computing time than the individual models. In conclusion, we selected EXT, RDF, MLP and the voting model to achieve the objective to classify large sets of ncRNA sequences accurately and time efficiently.

**Fig. 5.**
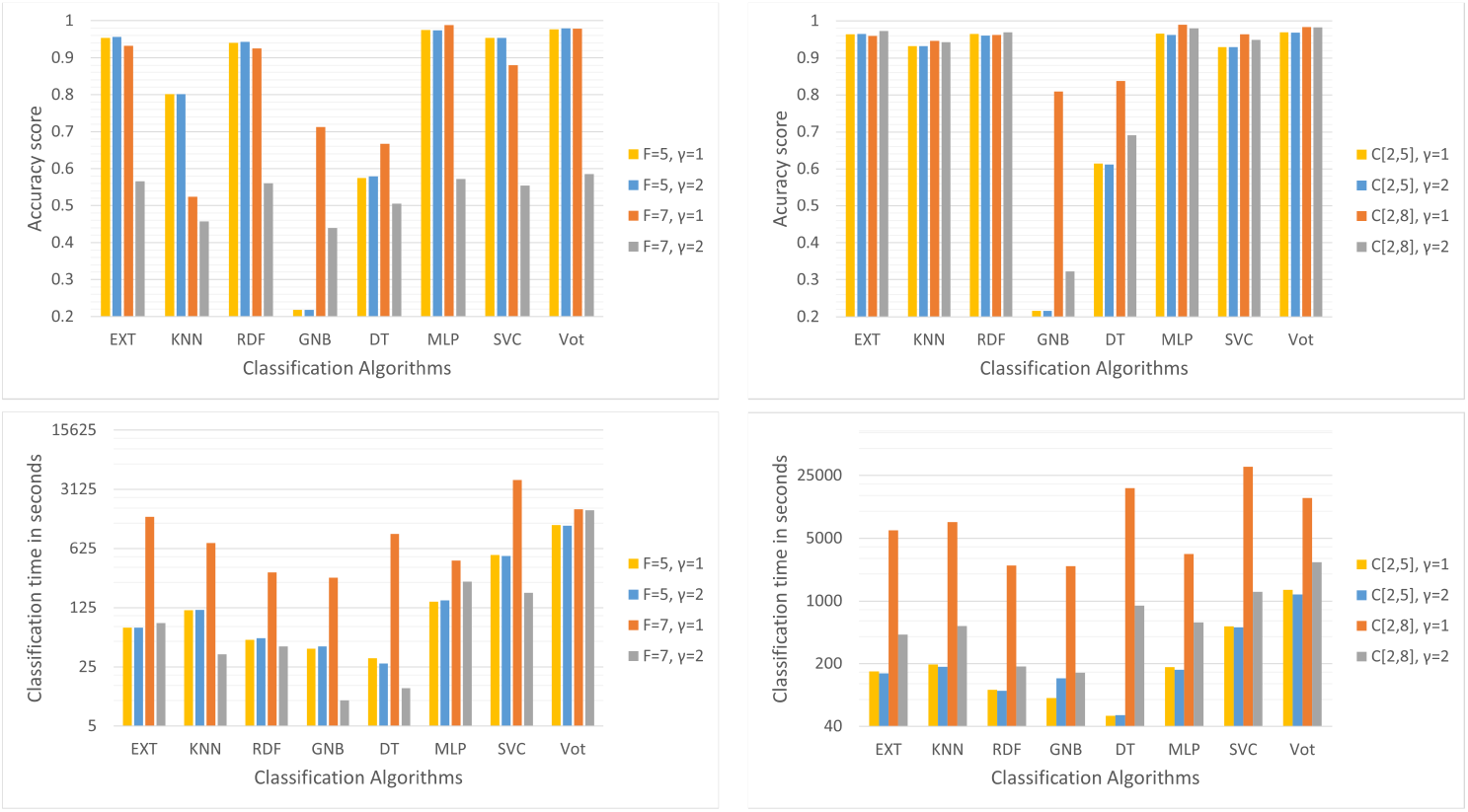
Classification accuracy (Top figures) and time (Bottom figures) for seven classification algorithms and a voting model under previously selected parameter settings for fixed-length motifs (Right figures) and combined-length motifs (Left figures).

### 3.3 Methods comparison

In this section, we describe and discuss the results of the experiments conducted to compare the different versions of our method with existing sequence and structure-based classification methods on the three ncRNA family datasets.

#### 3.3.1 Comparison EXT, RDF, and MLP on Dataset #1

Figure 6 and Supplementary Figure 6 show the test scores (accuracy and F-beta score) of the classifications obtained using EXT, RDF, MLP and the voting model after training the models on the whole training dataset, with the different parameter settings selected: either fixed length *F* = 5 or *F* = 7, or combined length motifs *C*[2, 5], *C*[2, 7] or *C*[2, 8]; *α* = 50; *γ ∈ {*1, 2*}*. Regarding the classification accuracy, the test scores were correlated to the validation scores obtained in the feature and model selection step on the medium dataset. The test scores are lower but remain high, with most of them larger than 0.85. For the classifier EXT, the setting (*C*[2, 7]; *γ* = 2) yields the highest accuracy, and for RDF, its (*C*[2, 7]; *γ* = 1). Unlike EXT and RDF, MLP yields better results with the fixed length motifs than with the combined length motifs. For MLP, the setting (*F* = 7; *γ* = 1) yields the highest accuracy. As expected, the voting model improves the accuracy in the four models. Regarding the F-beta scores, they are lower than the accuracy scores because of the dataset imbalanced property. Yet, both scores are correlated, and the F-beta scores are also high, which support comparisons results based on the accuracy scores. The computing time for training and testing is also an important criterion for the comparison. The test times of the three classifiers are comparable, but the training times differ. The MLP classifier takes about one hour in average, while EXT and RDF require about 3.5 hours each in average. The reported computing times average all parameter settings, but the experiments with fixed length motif datasets require less time than combined length motif datasets because of the datasets smallest size.

**Fig. 6.**
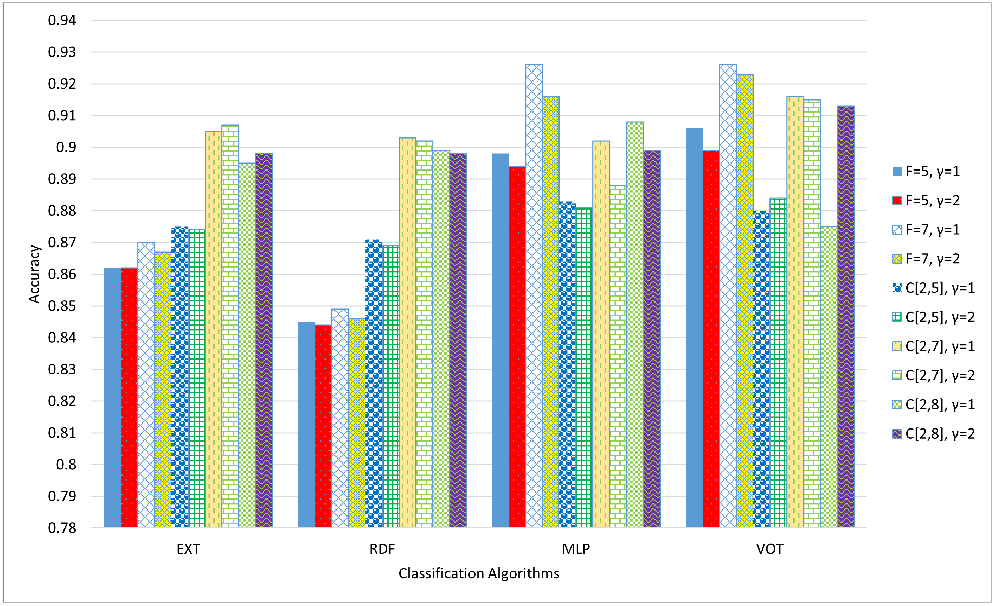
Test accuracy score for EXT, RDF, MLP and the voting model on Dataset #1.

#### 3.3.2 Comparison with Infernal, Blast and DeepNcRNA using Datasets #1 and #2

This section compares the different versions of our method with three existing sequence or structure-based classification methods on Datasets #1 and #2. The compared methods are Infernal (Nawrocki and Eddy, 2013), deepncRNA (Noviello et al., 2020), and Blast (Altschul et al., 1990). For Infernal, we included three versions: the default sequence- and structure-based version that uses HMM filters and CMs denoted by Inf_hmm_cm, the sequence-based version that uses only HMM filters denoted by Inf_hmm, and the structure-based version that uses only CMs denoted by Inf_cm. For Blast, we created a database using the training dataset. Then, for each sequence in the test dataset, we performed a Blast search using word size = 7. We tested various classification methods relying on Blast results (data not shown), but here, we report only the best performing method results denoted by Blast(av-s). Given the results of a Blast search for a test sequence, Blast hits in the training dataset were grouped by ncRNA family, and for each family, we computed an average raw similarity score. The test sequence was assigned to the family having the highest score. DeepncRNA is a sequence-based deep learning method. The best performing version of deepncRNA represents each sequence as a binary matrix such that columns represent k-mers (e.g., 16 columns for 2-mers) and rows represent the consecutive non-overlapping k-mers of the sequence (e.g., for a sequence of length 100, there are 50 non-overlapping consecutive 2-mers). The cell (*i, j*) of the matrix receives the value of 1 if the *i*^th^ k-mer of the sequence is *j*^th^ k-mer in the list of k-mers, otherwise it receives the value of 0. Using this binary matrix representation of RNA sequences, deepncRNA achieves the classification using a Convolutional Neural Network (CNN) composed of 3 layers of nodes. Depending on the length of k-mer considered in the experiments, the method is denoted by deepncRNA1 for 1-mer, deepncRNA2 for 2-mer, or deepncRNA3 for 3-mer.

Figure 7 and Supplementary Figure 15 show the results of the comparison. On Dataset # 2, our method (MLP version) yields the highest accuracy scores with the lowest classification times for all noise percentages. The method is also robust to the addition of noise. Infernal methods rank second in terms of classification accuracy, but the method performance decreases rapidly when we add noise. The Infernal methods also require huge model generation times and classification times. Blast ranks third in terms of classification accuracy, and is also robust to the addition of noise. While the classification time of Blast is high and comparable to that of Infernal, its model generation time is very low compared to the other methods. The deepncRNA methods rank last in terms of classification accuracy, but they are also robust to the addition of noise. Note that the results reported for deepncRNA are from (Noviello et al., 2020) because the implementation provided with the method did not work. This is why we did not report the model generation time and the classification time for deepncRNA. On Dataset #1 (Supplementary Figure 15), the comparison was limited to 400 families because of the high model construction time required by Infernal. Also, the comparison left deepncRNA out because of the lack of a working implementation. The results were like those obtained on Dataset #2, however, this time, Infernal outperforms other methods in terms of classification accuracy, closely followed by Blast and the MLP version of our method.

**Fig. 7.**
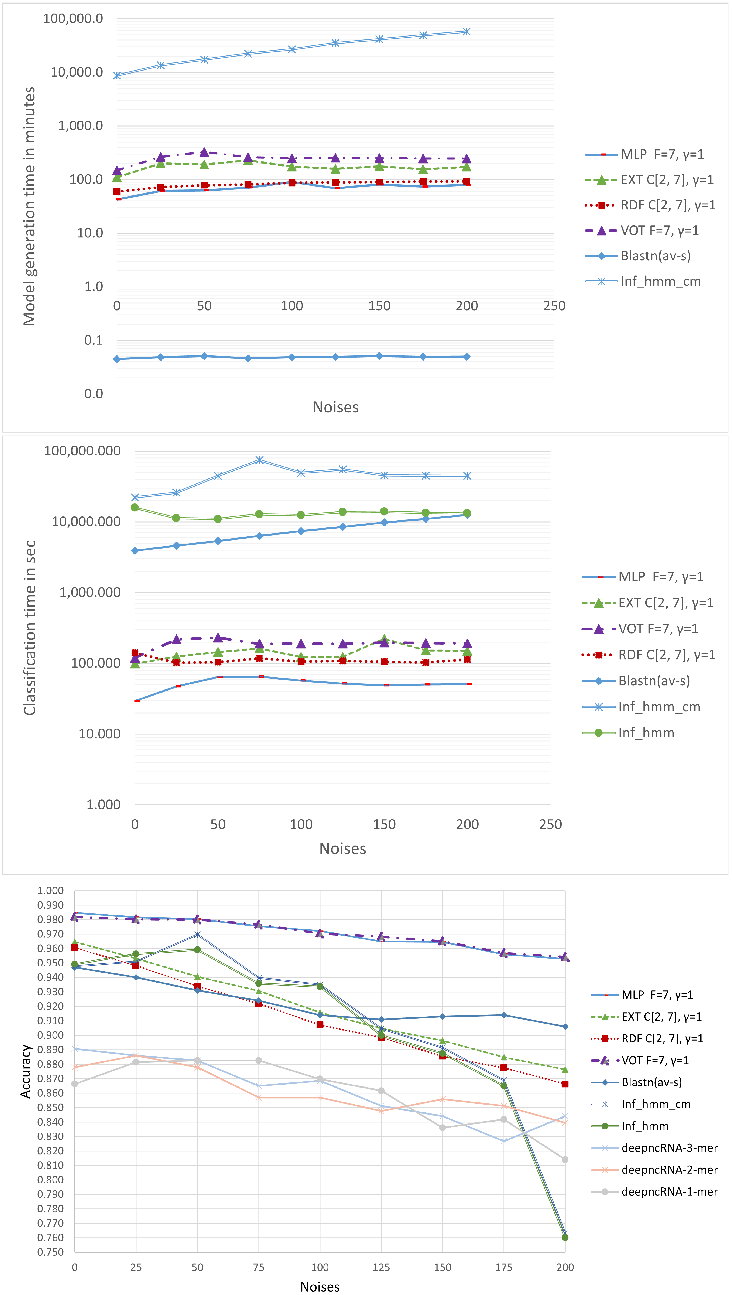
Model generation time, classification time and accuracy of compared methods on Dataset #2.

In conclusion, the sequence- and structure-based methods of Infernal achieve a very good classification accuracy at the cost of a huge computing time for model construction and classification. However, the performance of Infernal methods varies depending on noise. On the other hand, our methods and Blast, which are sequence-based methods, achieve a slightly similar accuracy, but with an extraordinarily lower time for model construction and classification, and a good robustness regarding noise. Our method, which uses a selection of common sequence motifs to represent RNA sequences, also outperforms the deepncRNA that relies on k-mers for RNA sequence representation.

#### 3.3.3 Comparison with Infernal and Blast using Dataset #3

Figure 8 shows the classification accuracy of the methods on the low sequence similarity and the high sequence similarity datasets of Dataset #3. For high sequence similarity within RNA classes, all methods perform very well. Infernal methods perform particularly well on this dataset. However, for low sequence similarity within classes, Infernal methods perform poorly, especially with HMM filters. That CM only performs better with HMM filters is not surprising; CM-based classification uses information on the conserved secondary structure, while HMMs rely on a globally conserved sequence which this dataset lacks. On the other hand, our methods and Blast perform very well on this dataset.

**Fig. 8.**
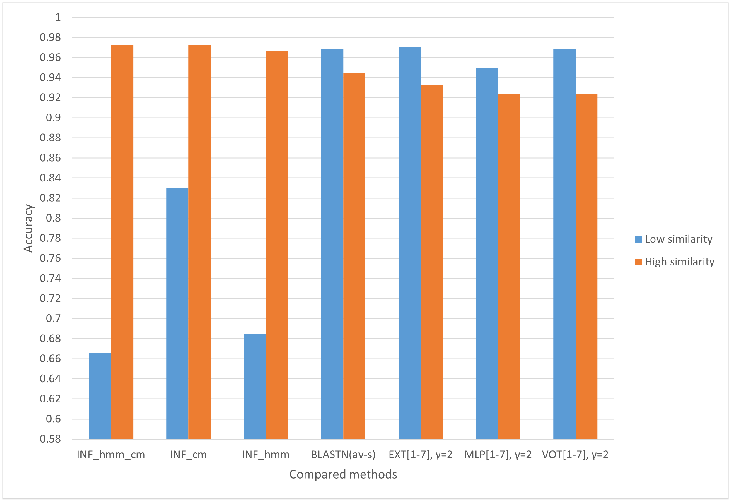
Classification accuracy of compared methods on Dataset #3.

We conclude that when the global sequence similarity is low within RNA classes, methods that rely on local sequence motifs perform better, like our method or Blast, than methods that rely on global sequence or structure similarity. Using local sequence motifs even outperforms the use of secondary structure information in this case.

## 4 Conclusion

Classifying ncRNA sequences is an important task for genome annotation and RNA functional analyses. We need efficient methods to effectively perform the classification of the large amounts of new ncRNA sequences discovered by high-throughput RNA sequencing. In this context, machine learning (ML) approaches are expected to provide efficient solutions.

Using the appropriate features to represent the data is crucial to achieve an efficient and effective classification using ML classifiers. This paper introduces a novel sequence-based classification method that uses a supervised learning approach. Unlike current ML classification methods that typically rely on k-mer and/or secondary structure features, our method relies on the computation and the selection of common sequence motifs to classify ncRNAs. This approach yields a fast and accurate classification.

The results of the method suggest that relying on locally conserved sequence motifs performs better than relying on k-mer or secondary structure based representations. Compared to existing methods, our method achieve a similar or higher accuracy with an extremely lower computing time for classification, and a good robustness with respect to noise.

However, the fact that our method relies solely on sequence information is both an advantage and a limitation. It avoids the huge computation time associated with using secondary structure information, but it lacks structure conservation information that could improve the classification. Future work will explore ways to improve the classification by integrating information about the structure without affecting the efficiency of the method. This could be achieved by adding local structural motifs such as super-n-motifs (Glouzon et al., 2017) to represent RNA sequence and structure.

Future extensions of the method may be to use the models to search sequence databases like whole genomes for members of the RNA families included in the training data. This will require to design and to include noise models in the method. The preliminary results obtained on the robustness of the method with regards to noise suggests that it will also enable a fast and accurate annotation of genomes. Another direction is to use the models to predict novel unknown ncRNA families. Based on the method’s ability to accurately classify ncRNA clans, this could be achieved by identifying sequences of unknown ncRNA families that are included in known ncRNA clans.

## Acknowledgements

We thank Shengrui Wang (CS Department, Université de Sherbrooke) for his insights and clever advices on the experimental design of this study.

## Appendix A Supplementary figures

**Fig. 9.**
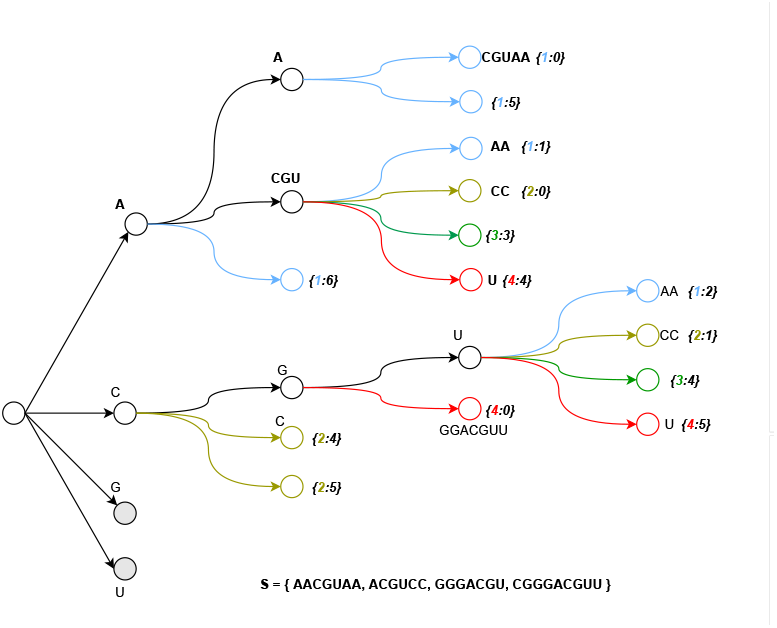
Partial view of the GST for S ={ AACGUAA, ACGUCC, GGGACGU, CGGGACGUU }. Only the subtrees starting with A or C are extended. Any substring *x* that is present in at least one of the RNA sequences is represented only once in the GST by a path from the root to a node *n*_x_. The set of leaves of the subtree rooted at *n*_x_ are labeled with information on all occurrences of substring *x* in the RNA sequences. Each leaf is labelled with the start location of the substring in an RNA sequence, and the corresponding RNA sequence identifier. For instance, substring ‘ACGU’ is present at location 1 in sequence 1, at location 0 in sequence 2, at location 3 in sequence 3, and at location 4 in sequence 4.

**Fig. 10.**
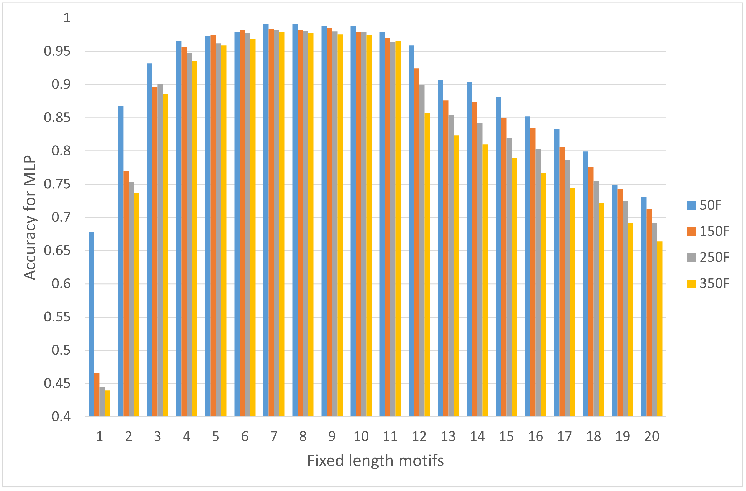
Relation between the size of fixed-length motifs and the classification accuracy obtained using MLP.

**Fig. 11.**
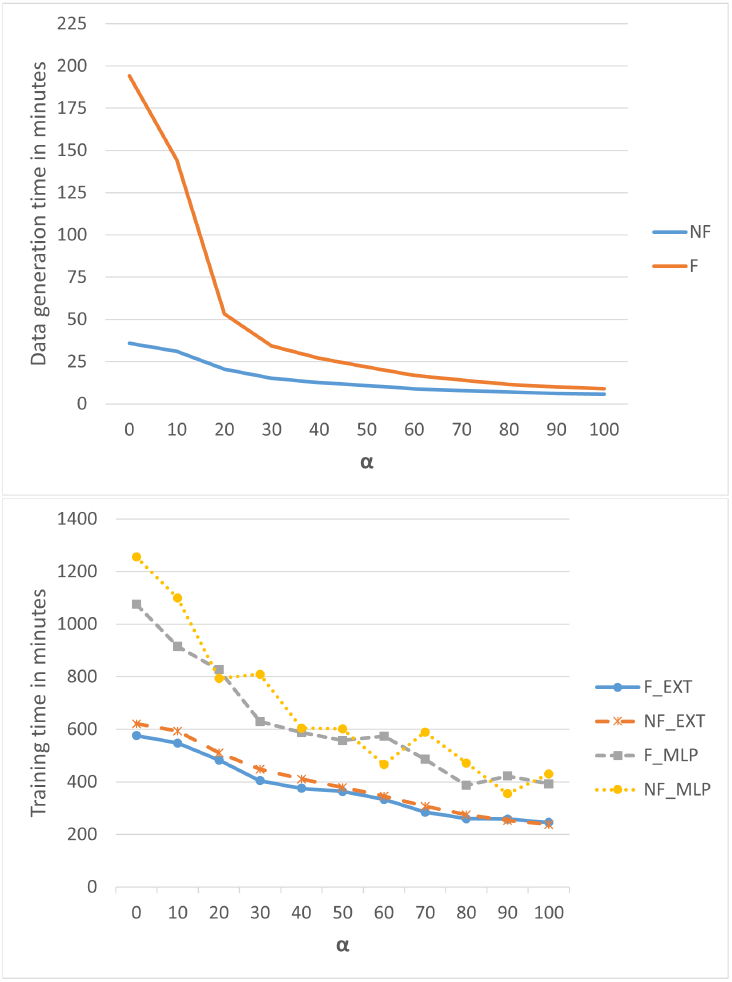
Relation between *α* and : (Top figure) the data generation time; (Bottom figure) the training time for EXT and MLP, using *submotif* = F or *submotif* = NF.

**Fig. 12.**
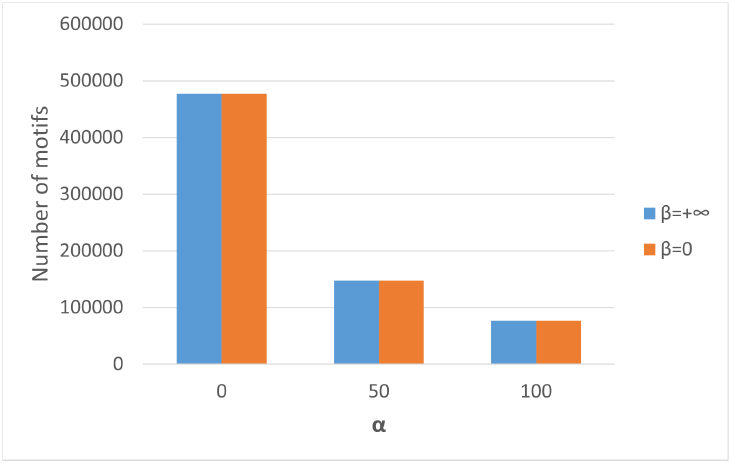
Comparison of the numbers of motifs for *β* = 0 and *β* = +*∞* for different values of *α*.

**Fig. 13.**
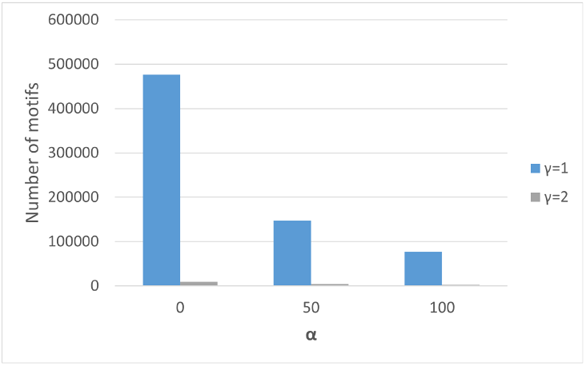
Relation between *α, γ* and the number of generated motifs.

**Fig. 14.**
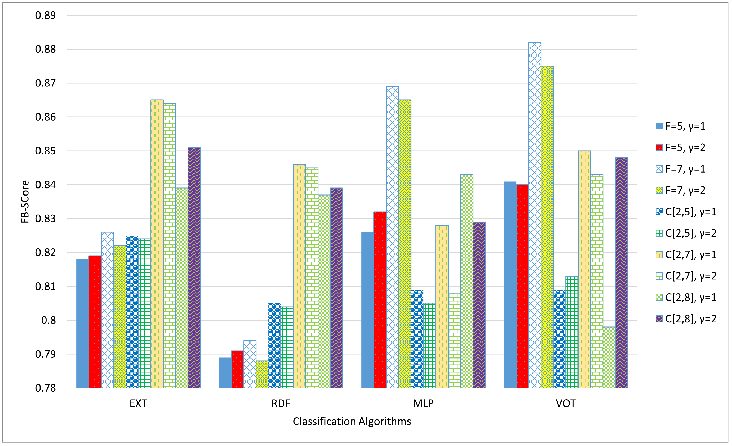
F-beta score of the classification for EXT, RDF, MLP and the voting model on Dataset #1.

**Fig. 15.**
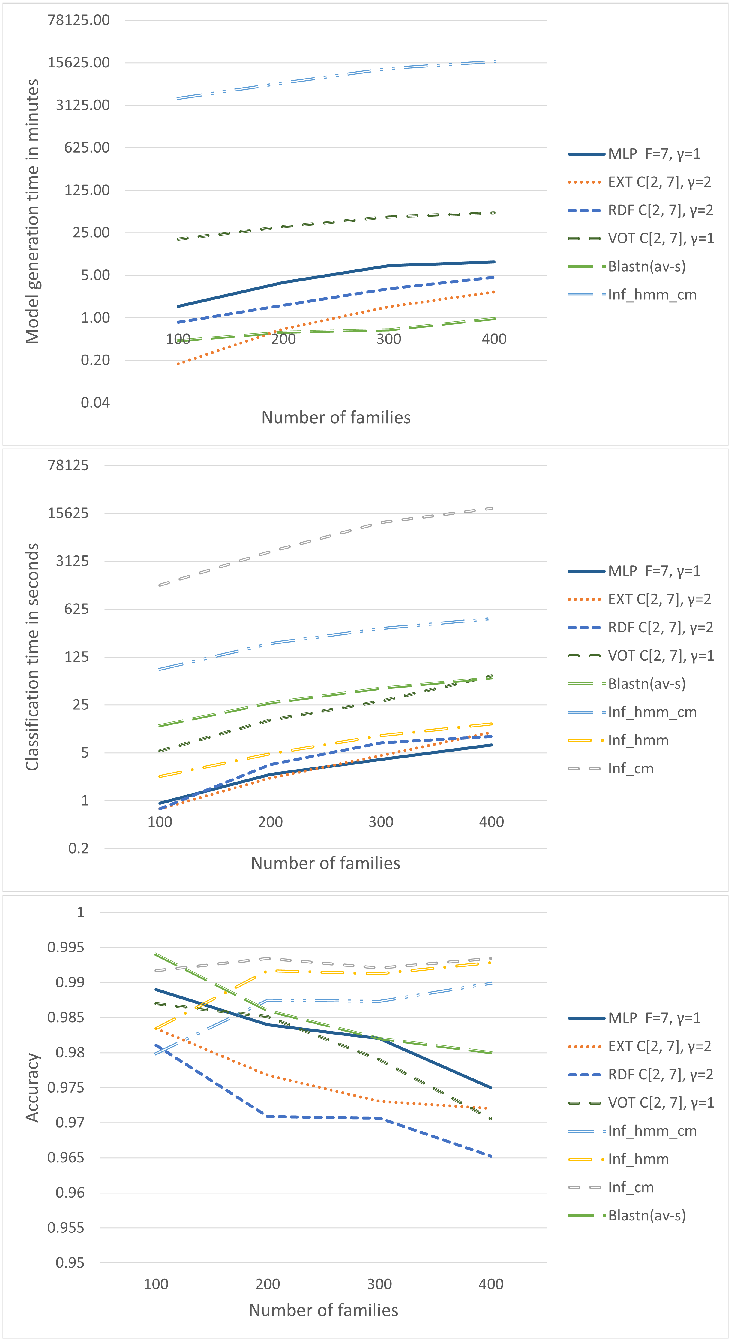
Model generation time, classification time and accuracy of compared methods on Dataset #1.

